# AInsectID Version 1.1: an Insect Species Identification Software Based on the Transfer Learning of Deep Convolutional Neural Networks

**DOI:** 10.1101/2024.11.01.621497

**Authors:** Haleema Sadia, Parvez Alam

## Abstract

AInsectID Version 1.1^1^, is a GUI operable open-source insect species identification, color processing^2^ and image analysis software. The software has a current database of 150 insects and integrates Artificial Intelligence (AI) approaches to streamline the process of species identification, with a focus on addressing the prediction challenges posed by insect mimics. This paper presents the methods of algorithmic development, coupled to rigorous machine training used to enable high levels of validation accuracy. Our work integrates the transfer learning of prominent convolutional neural network (CNN) architectures, including VGG16, GoogLeNet, InceptionV3, MobileNetV2, ResNet50, and ResNet101. Here, we employ both fine tuning and hyperparameter optimization approaches to improve prediction performance. After extensive computational experimentation, ResNet101 is evidenced as being the most effective CNN model, achieving a validation accuracy of 99.65%. The dataset utilized for training AInsectID is sourced from the National Museum of Scotland (NMS), the Natural History Museum (NHM) London and open source insect species datasets from Zenodo (CERN’s Data Center), ensuring a diverse and comprehensive collection of insect species.

## INTRODUCTION

There are several ways in which insect species can be identified. These include methods based on observed morphological characteristics ^3^, which in turn might require microscopy ^4^, direct comparison against insect collections and reference specimens ^5^, the identification of unique anatomical features through dissection ^6^, observation of specific behavioral characteristics ^7^, chemical analysis methods ^8^, geographic distribution data ^9^ and DNA barcoding ^10^. Each method can be used either individually, or, in parallel with other identification methods to reduce uncertainty levels. These techniques often require a high level of expertise to ensure that identification is accurate. This is especially true when identifying insects with intra-class variations, which are generally more challenging to identify as they may exhibit several similar morphological traits, and may even share similar geographical locations. In the absence of experts or trained personnel, insect identification can be challenging ^11^, especially if instant identification is required during field-work expeditions, or if resources are limited. Collaborative efforts alongside the use of advanced techniques like DNA barcoding ^10^ can be useful to improve the precision in identification, which is a further testimony to that the process of insect identification can be laborious, time-consuming, resource-extensive, and often also requires communication with experts on specific types of insect. The rapid growth of artificial intelligence (AI) provides opportunities for advancements in this area ^12^, and if built accurately, may enable the rapid identification of insect species without the need for specialty resources or skills. Common AI models employ machine learning (ML), or deep learning (DL) algorithms, which are themselves subdivided into different techniques. We focus on two specific techniques in this paper to develop insect identification algorithms. These include convolutional neural networks (CNN) and transfer learning (TL), and we will discuss each of these in more detail in the following two sections, with reference to previous work where these techniques have been used for species identification.

### Convolutional Neural Networks and Species Identification

CNNs are popular deep learning architectures ^13^ well-suited for insect identification as they automatically learn complex, hierarchical representations from images and effectively handle object detection and classification tasks ^14 15^. There are numerous examples of CNN based models that have recently been developed for species identification purposes. CNN-based object detection models for example, such as Faster R-CNN and YOLO, can locate and identify nine dominant insect species across more than two million images with a 93.80% accuracy within video streams ^16^. The DL cloud-based platform, Deep Automated Bee Identification System (DeepABIS), consists of 9942 images, 166 genera, and 881 species. These are used to identify bees showing a 93.95% accuracy ^17^. Motta et al. ^18^ trained CNN to automate the morphological classification of mosquitoes, collecting a dataset of 4056 mosquito images of three species: *Aedes aegypti, Aedes albopictus*, and *Culex quinquefasciatus*, and evaluated the images against three different neural network architectures, LeNet, AlexNet, and GoogleNet, achieving 57.5%, 74.7% and 83.9% levels of accuracy, respectively. Their work shows us that network architecture selection is a critical part of the process, if high performance models are to be developed for species identification. Hoyal et al. ^19^ focused on quantifying phenotypic similarity by calculating (Euclidean) phenotypic distances between interspecies co-mimics using deep convolutional neural networks (DCNN), achieving an 86% accuracy from a dataset of 2468 butterfly images covering 38 subspecies of *Heliconius erato* and *Heliconius melpomene*.

Achieving high accuracy in identifying intra-class insect variants is challenging for CNN models. While DL algorithms perform well in species classification, training is complex due to the need for high-resolution, detailed, and labeled datasets. Insect datasets often face class imbalances, with limited data available for rare or newly discovered species, complicating CNN training. Additionally, insects photographed in natural environments introduce further challenges due to intricate backgrounds and details. Insects may be photographed from various angles and postures, with wings and limbs in different positions. Additionally, variations in color and texture caused by lighting fluctuations complicate feature extraction ^20 21^. To address challenges in deep learning, researchers have been exploring techniques such as transfer learning, few-shot learning, and data augmentation to enhance the data efficiency and adaptability of algorithms for various tasks and scenarios ^22^. The following section focuses on transfer learning as it is of direct relevance to how we have developed prediction algorithms in AInsectID Version 1.1.

### Transfer Learning and Species Identification

TL in DCNN is a way of leveraging knowledge gained while training a model on one task, and subsequently applying the knowledge to a different but related task ^22 23^. The idea is to transfer the learned representations of features from one domain to another, usually from a pre-trained model on a large dataset (such as ImageNet Dataset) to a new, small, and different but related dataset, Figure 1. Fine-tuned TL is especially beneficial when dealing with small datasets or when retraining from scratch is impractical due to computational or time constraints. This approach involves further training a pre-trained model on new data by unfreezing selected layers, allowing updates based on the new dataset. Careful consideration of pre-trained CNN model selection, the extent of fine-tuning, data augmentation strategies, and the evaluation of model performance on a new dataset is essential ^24^. When fine-tuning only a subset of layers, the layers excluded from the process are said to be “frozen.” This technique enables the refinement of higher-level feature representations within the base model. Recent studies have introduced diverse methods with distinct strategies to enhance conventional fine-tuning ^25 23^. Commonly employed strategies following this approach involve either fine-tuning all layers of the neural network, as detailed in ^26^, or selectively fine-tuning only the last few layers, as demonstrated in ^27 28^. Hyperparameter optimization is important as it involves the systematic exploration of optimal hyperparameter sets to enhance model performance. Both fine-tuning and hyperparameter optimization are important in achieving optimal performance in DL models ^29 30 31 32^.

**Figure 1:**
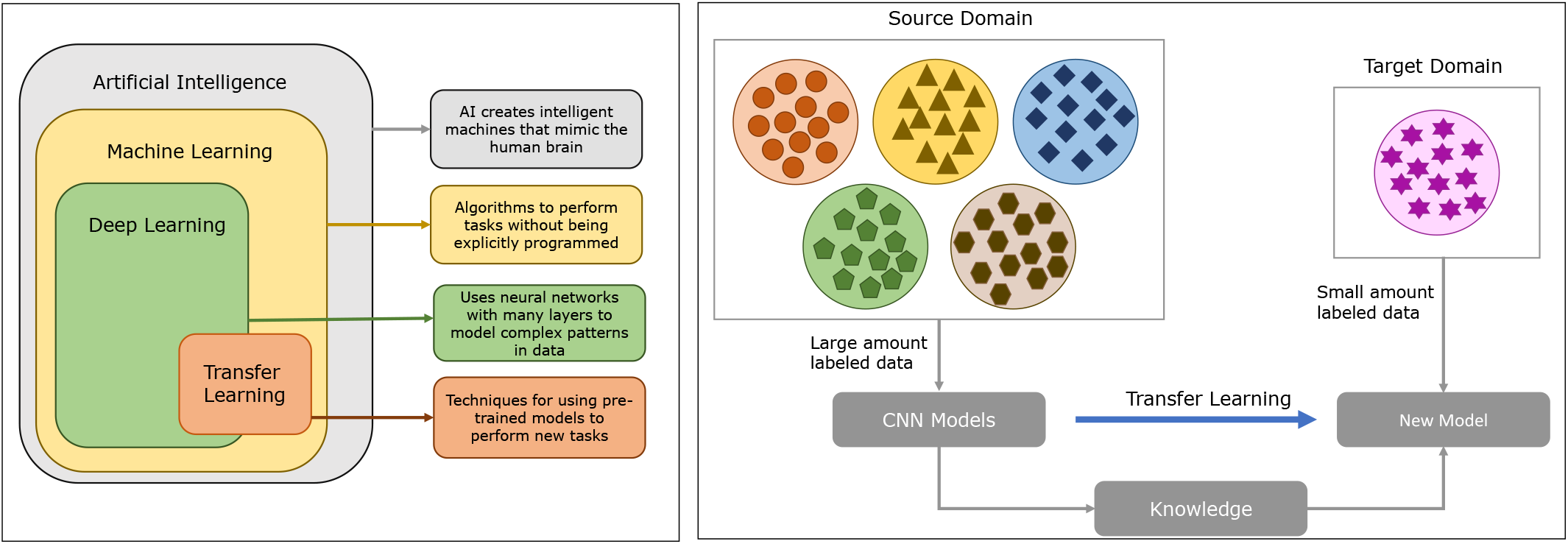
(Left) Visual Representation of Relationship between Artificial Intelligence, Machine Learning, Deep Learning, and Transfer Learning. (Right) Illustration of Transfer Learning

Examples of both fine tuning and hyperparameter optimization have been tasked with species identification. ResNet for example has been used to classify forest insects, using a pre-trained model and fine-tuning for 30 insect species, achieving a 94% classification accuracy ^33^. Using TL to reduce training time is an effective strategy in deep learning, especially when dealing with limited computational resources or small datasets. Fathimathul et al. ^34^ for example, trained various CNN models including VGG16, VGG19, MobileNet, Xception, ResNet50, and InceptionV3 by TL to identify butterfly species. They collected 10035 images from 75 different species of butterflies using the Kaggle website and report that the InceptionV3 architecture outperformed all other architectures, achieving an accuracy of 94.66%. Yasmin et al. ^35^ developed an android application to classify 10 individual butterfly species consisting of 832 images empploying three different methodologies including, principal components analysis (PCA) with a support vector machine (SVM), DL by 4-Conv CNN model, and TL using a pre-trained VGG19 architecture. The most successful of these was through TL using a pre-trained VGG19 model achieving a 96.5% accuracy. Shamim et al. ^36^ identified vector and non-vector mosquito species at an accuracy of 97.02% comparing DCNN models VGG-16, Inception V3, and MobileNetV2 assisted by TL. Agarwal et al. ^37^ identified 102 pest species by collecting a data set consisting of 12956 total images including 80 species of 12 genera. They trained five DCNN models (VGG-19, ResNet-34, ResNet-50, ResNet-101, and DenseNet-121, DenseNet-169, MobileNetV2, ResNet50), the pretrained VGG-19 showing significant success with a 93.47% accuracy. Soni et al. ^38^ designed crop pest detection mobile applications by training four DCNN models (Xception, MobileNet, MobileNetV2, and EfficientNetB7), the highest accuracy of which was delivered by EfficientNetB7 at 95.6%. There are several apps that help to identify insect species typically using image recognition technology to match an image of the insect with a database of known species. These include: iNaturalist, a community-driven using crowd sourced identification and expert verification ^39^, similar in essence to BugGuide.net ^40^, PlantSnap, which is primarily designed for identifying toxic plants but can also identify many insects ^41^, and a forest insect classification app developed by Lim et al. ^42^ to classify 30 forest insects using ResNet with an accuracy of 94%. While these apps can be very useful their outputs often require cross-referencing against additional sources and expert opinions, also generally requiring high quality input images to enable identification accuracy ^43^. Fundamentally, the implementation of TL comes with its challenges ^23^, which have gained increased attention in recent times ^44 45 46^. To effectively customize a pre-trained model for a new task, certain components (layers) require retraining, while others must remain unaltered. Deciding which layers should be activated for training (fine-tuning) and which ones should remain frozen, is a necessary challenge in the adaptation process. Moreover, high dimensionality, computational cost, non-convexity, limited domain knowledge, interactions, data sensitivity, black-box nature, noisy evaluation, and the risk of overfitting to validation sets pose additional challenges in hyperparameter optimization ^47^.

### Insect Mimicry - an Additional Complexity

More than 50% of the Earth’s biological diversity are represented by insects ^48^, with over 1.02 million species having been described to date and an estimated 80% yet to be discovered ^49^. Within single species, insects can exhibit considerable intra-species diversity ^50^, which can increase the challenges associated with insect identification. Differentiating between insects at higher taxonomic levels (order and family) is generally more straightforward than at lower taxonomic levels (genus and species). Morphological distinctiveness is wide ranging at higher taxonomic levels and features at this level are often easier to identify, making it possible to position insects into broader categories. Insect identification becomes more challenging when descending the taxonomic hierarchy. This is due to increased variability and complexity among closely related species at the lower taxonomic levels. Moreover, many insect species have intra-class variations that reflects insect adaptation broad ranging ecological conditions such as geographical location, habitat, food source, environmental conditions, and evolutionary history ^51 16 33^. A fascinating adaptation strategy in the insect world is mimicry. Insect mimics have evolved to resemble other species in their environment. This serves various purposes from protection to resource security, Figure 2. Insect mimicry includes visual appearance, behavior, and sometimes even chemical cues ^51^.

**Figure 2:**
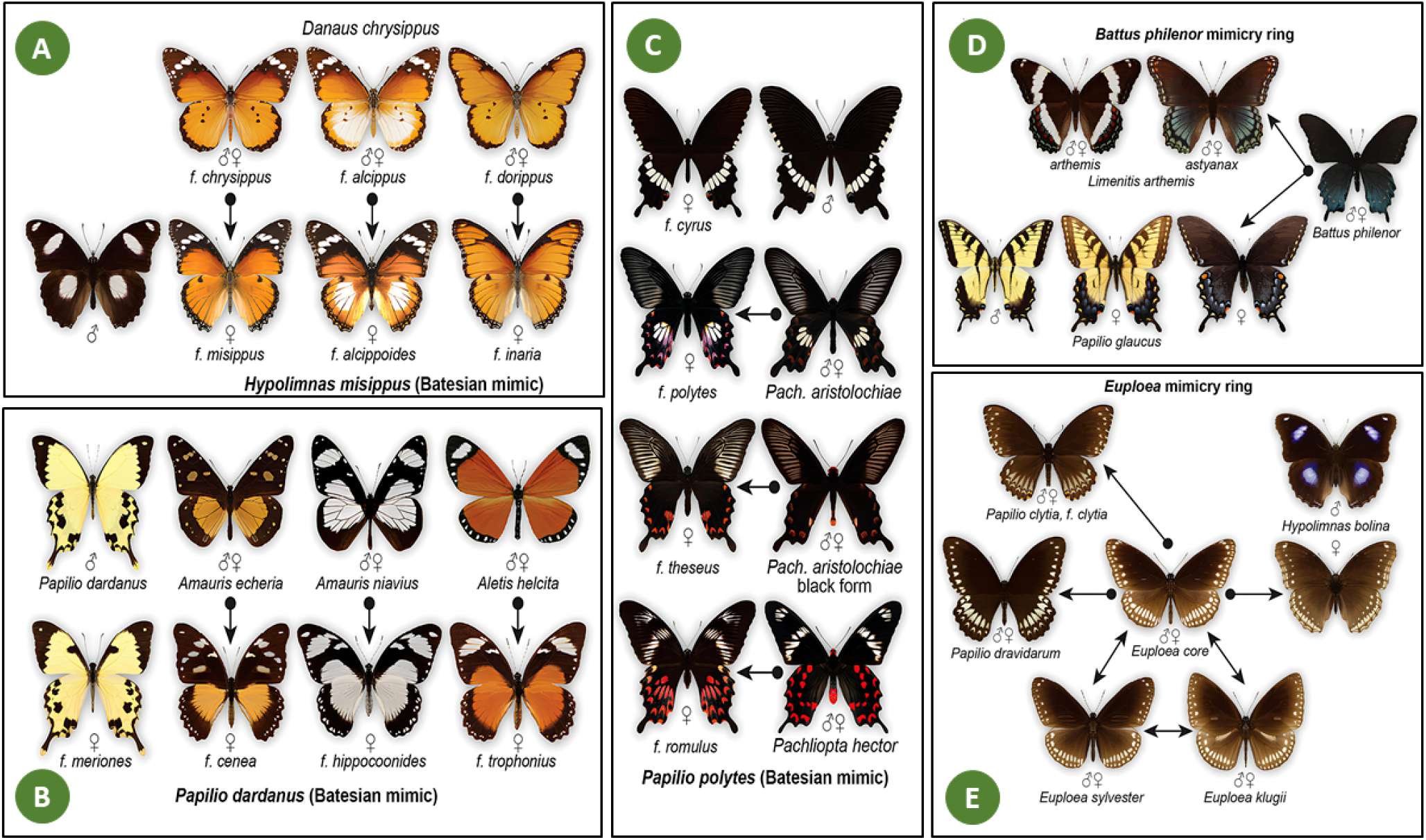
(A) *Danaus-Hypolimnas* mimicry ring illustrates a rare example where both the Batesian model and the mimic are polymorphic, female forms of the Batesian mimic *H. missipus* mimicking *D. chrysippus* in a form-specific manner. (B) *P. dardanus*, and (C) *P. polytes*, multiple female forms mimic distinct species of models, the male and male-like female forms are nonmimetic, which in *P. polytes* also represents the ancestral phenotype, represent the degree to which natural selectionthrough predationmay drive nearly perfect and polymorphic wing pattern resemblance between Batesian models and mimics. (D) and (E) are two mimicry rings are driven by the aposematic species, *B. philenor* of North America and *Euploea* of S. Asia respectively, which are mimicked by multiple Batesian mimics. Figures reused from ^52^ with the permission of Wiley Periodicals, Inc.

In this paper, we use VGGNet, GoogLeNet, InceptionV3, MobileNetv2, ResNet50, and ResNet101, models to train species datasets with the primary goal of achieving high prediction accuracy. Our focus is to enable precise differentiation between mimic species and their mimicked counterparts, ensuring the models can effectively distinguish subtle species-specific features. We choose these models as baselines, since each has reported evidence of high image-based prediction accuracy rates. In terms of insect dataset classification through transfer learning, this work will investigate the impact of various optimization algorithms on selected pre-trained CNN architectures. We will explore transfer learning scenarios in conjunction with four optimization algorithms: SGD, SGD with momentum, RMSProp, and ADAM, to identify the optimal hyperparameters. Data augmentation techniques will finally be employed to address any challenges related to both class imbalance and limited data (in the cases of certain rare species).

## METHODS

### Data Collection and Preprocessing

Deep learning models can automatically learn to distinguish relevant features in image datasets during training. With a large and diverse dataset, these models uncover complex patterns, enhancing identification accuracy and model generalization while reducing the risk of overfitting. ^53^. Taking these points into consideration, here, a total of 40,000 images from 150 insect species (butterflies, moths, bees, and dragonflies) including mimics were collected to form the datasets. Our datasets comprised 22,000 images with white backgrounds and 18,000 images with complex/colored backgrounds. The dataset containing images with a white background was collected from the National Museum of Scotland (NMS), Edinburgh, Scotland, and the Natural History Museum (NHM), London ^54^. The dataset containing images with a colored background was collected from the open-source dataset of insect species from Zenodo ^55^, an open science repository hosted by CERN’s Data Center. The datasets were properly labeled by creating individual repositories for each insect species, with each repository assigned a unique class label following the form, “Species_name”. Images were then organized within these species-specific folders. Our dataset includes five mimics of *Danaus plexippus*, two mimics of *Delias belisama*, and two mimics of *Battus philenor*. Our training process of the Deep Convoluted Neural Networks (DCNNs) developed in this work is outlined in Figure 3.

**Figure 3:**
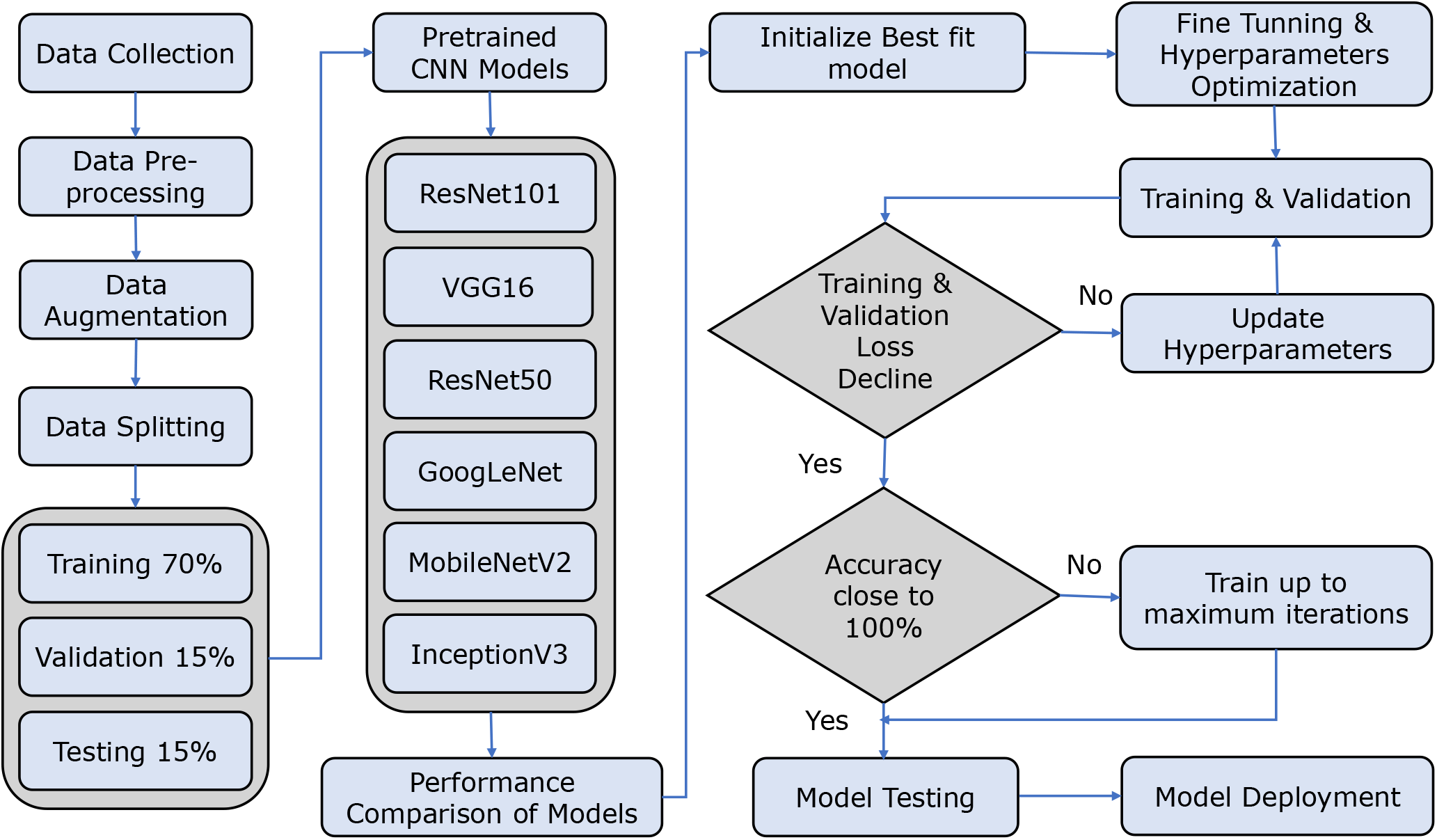
Flowchart outlining the training process of Deep Convolutional Neural Networks (DCNNs) for insect species identification. The chart begins with data collection and preprocessing, followed by data augmentation to enhance diversity. The dataset is then split into training, validation, and testing sets. Several pre-trained models (VGGNet, GoogLeNet, InceptionV3, MobileNetV2, ResNet50, and ResNet101) are evaluated for accuracy, and the top-performing model is selected for further training. This model undergoes an iterative fine-tuning process, with continuous parameter adjustments to enhance performance until the desired accuracy is reached, facilitating the precise differentiation between mimic species and their mimicked counterparts.

#### Data Augmentation

Data augmentation is the process of artificially generating new data from an existing dataset, effectively increasing the available training data and improving performance in image classification tasks ^53 56 57^. A challenge we faced was that training data for many of the rarer insect species were sparse, time-consuming, and resource-intensive. We applied data augmentation techniques including random rotations, scaling, flipping, shearing, translating, cropping, and brightness adjustments. We initially cropped each image into four sections, after which we applied other data augmentation techniques such as scaling and flipping the cropped images. The process of image cropping results in a 4-fold increase in training data. Additionally, horizontal and vertical flipping was used alongside random scaling, shearing, and translation processes, to further augment the training for species where the datasets were limited. We finally applied the shuffling technique, in which the order of images is randomly rearranged within the dataset to increase further variability during training, thereby preventing the model from learning spurious patterns based on the order of the data.

As shown in Figure 4, the outcome of the distinct augmentation method to the original images, thereby increasing dataset variability and diversity. Images were translated images along the x and y Cartesian axes, while brightness adjustments were made to enable model adaptation to the varying lighting conditions often encountered during fieldwork. Furthermore, randomly rotated images, aid the model in learning diverse orientations, which is crucial when dealing with species in diverse habitats. Cropping is essential for eliminating irrelevant background noise, particularly beneficial for species with intricate features. Flipped images enhance the model’s ability to recognize invariant features across orientations, while sheared images add variability. By incorporating these augmentation techniques, We increased our dataset from 42,000 samples to 610,000 samples, thus significantly enhancing the diversity and robustness of the dataset. To maintain consistency, we implemented specific pre-processing steps to resize the input images in the dataset to 224 × 224, aligning with the input size requirements of the DL model. The dataset was split into three subsets: 70% training, 15% validation, and 15% testing for training DCNNs. Each adapted DCNN model was trained on the training dataset with validation on a validation dataset, and model performance was evaluated on the test dataset. The test dataset is not seen by the model during training, ensuring an unbiased evaluation of its performance on new, unseen data.

**Figure 4:**
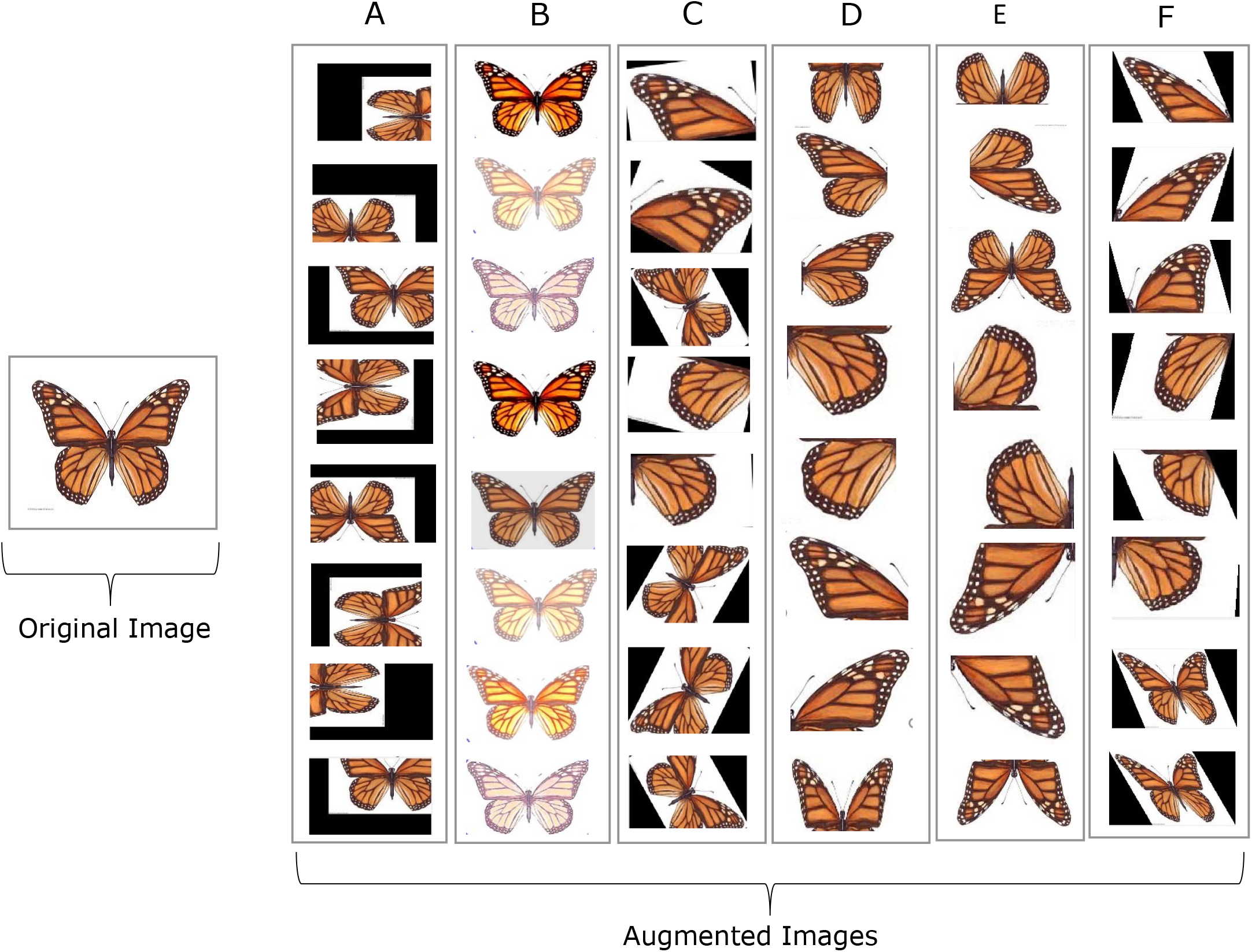
The figure shows the outcomes of applying various data augmentation techniques to images, including (A) Translation, (B) Brightness Adjustment, (C) Random Rotation, (D) Cropping, (E) Flipping, and (F) Shearing.

### Optimal DCNN Model Selection via Transfer Learning

TL was employed for DCNN model selection from renowned DCNN architectures, namely VGG16, GoogLeNet, InceptionV3, MobileNetV2, ResNet50, and ResNet101. We introduced additional layers on top of the pre-trained models with all existing layers frozen as shown in Figure 5. CNN architectures are originally designed to classify 1000 classes of ImageNet dataset, we replaced the classification layers, including the output layer, to align with the 150 classes in the insect species dataset and froze all the convolutional layers to retain the pre-trained feature extraction while fine-tuning only the newly adopted layers. In CNN layer architectures, the layers can be divided into feature extraction layers and classification layers. Feature extraction layers include convolutional layers, activation layers (ReLU), and pooling layers. They are responsible for capturing and extracting complex features from the input images ^15 12^. Convolutional layers perform a mathematical operation known as convolution, where a small filter (also referred to as a kernel) is systematically applied across the input image to extract features. Each filter is essentially a small matrix of numbers (typically 3×3 or 5×5), and these numbers are the **weights**. For example, if the filter size is 3×3, it will have 9 weights. During forward propagation, the filter slides over the input image and performs element-wise multiplication between the filters weights and the corresponding pixels of the image. The sum of these multiplications produces a **feature map**. The process of computing the output *Y* of a *i*^*th*^ convolutional layer is represented by Equation 1, where *X* denotes input feature map, *W* ^(*i*)^ is the set of filters, and *b*^(*i*)^ represents the bias vector associated with the *i*^*th*^ convolutional layer that adjusts the output of each neuron, enabling the model to capture complex relationships in the data effectively.

**Figure 5:**
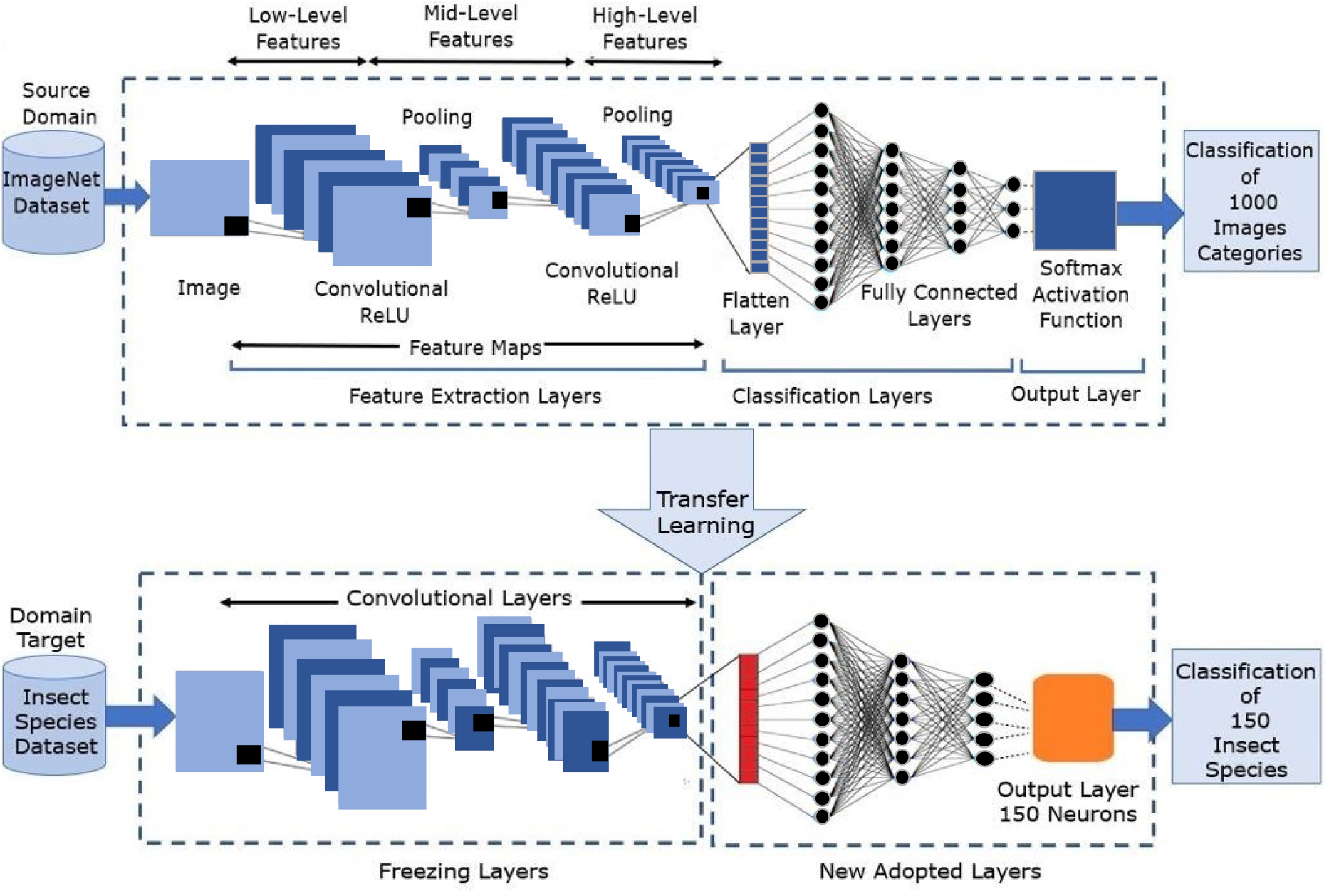
Illustration of our transfer learning pipeline that adapts ImageNet pre-trained Convolutional Neural Networks (CNNs) for insect species classification. The original CNN models are trained on 1,000 categories, enabling them to learn robust features. In this pipeline, new layers are added to classify 150 insect species while keeping the feature extraction layers of the pre-trained model frozen. This preserves the learned features from ImageNet, allowing the model to adapt to the specific characteristics of insect species. The training focuses on fine-tuning only the new layers, optimizing performance for accurate insect classification.

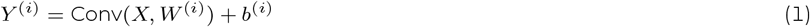

The activation layers introduce non-linearity, enabling network learning of complex patterns. The pooling layers downsample the feature maps, reducing their spatial dimensions while preserving relevant information. Low-level features including edges, textures, and shapes were presented by initial layers, while higher layers captured more abstract and complex features. High-level features are often the primary focus of fine-tuning, low-level features can still be relevant and can be adjusted ^58^. Once features are extracted, they are passed to the classification layers. Fully connected (dense) layers are frequently used for classification tasks. These take the flattened feature maps and map them to the output classes, using techniques such as softmax activation to produce probability scores for each class, indicating hence, the likelihood of the input’s association with each category ^15 12 16 17^..

After processing through preceding layers, the output layer generates logits for each class, which are then converted into probabilities using the softmax function for multi-class classification as shown in Equation 2, where *z*_*c* are the raw output scores for each class and *C* is the number of classes.

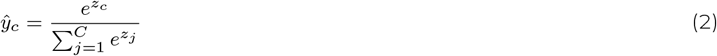

After obtaining the predicted probabilities, the categorical cross-entropy loss is computed by using Equation 3, where *y*_*c*_ is the true output label and 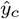 predicted output label.

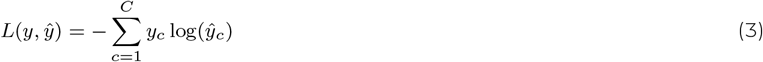

Backpropagation is used to compute the gradients of the loss function to each weight in the network as in Equation 4. This involves applying the chain rule to propagate gradients backward through the network layers. where 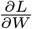 represents the gradient of the loss *L* concerning the weight *W*, 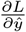 denotes the gradient of the loss for the predicted output 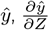 represents how the predicted output changes for the pre-activation value *Z*, and represents how the pre-activation value *Z* changes for the weights *W*.

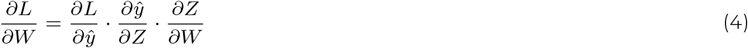

In the context of fine-tuning and unfreezing convolutional layers, this allows the model to update the weights and biases of the convolutional layers while keeping the weights of other layers frozen, as shown in Equation 5 and 6, where *α* representing the **learning rate**, is a small positive value, which controls the step size where the model parameters (weights and biases) are updated during training using gradient descent-based **optimization algorithms**. Stochastic Gradient Descent (SGD), SGDM (Stochastic Gradient Descent with Momentum), Adam, and RMSprop are optimization algorithms that were tested individually, at different learning rates ranging from 0.00001 to 0.01. ∇_*W*_ and ∇_*b*_ represent the gradients of the loss function *L* with respect to the weights and biases of the *i*^*th*^ convolutional layer, respectively.

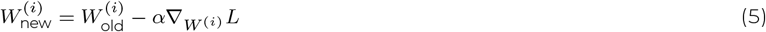

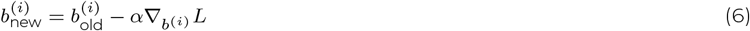

At each iteration *t*, the parameters *θ* are updated as defined by Equation 7. Here, ∇_*θ*_*L*(*θ*_*t*_) is the gradient of the objective function of *L*(*θ*) with respect to the parameters *θ* evaluated at *θ*_*t*_.

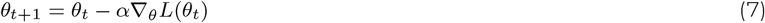

The optimization algorithm with momentum is based on Equation 8 and 9, where *β* is the momentum coefficient known as the decay factor that lies between 0 and 1, controlling the contribution of the previous momentum to the current update. The momentum term *v*_*t*_ accumulates the gradients over time with *β*.

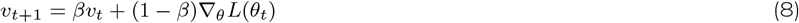

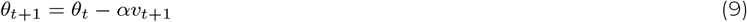

This experimentation aimed to identify the most suitable configuration that enhances the performance of the DCNN models on the insect dataset, and contributes valuable insights into selecting the best-fit pre-trained CNN model. By comparing the performance of all selected DCNN models, we identified the optimal pre-trained model to focus on fine-tuning and hyperparameter optimization to refine the selected DCNN model and push its performance closer to achieving 100% accuracy.

### Advancing Optimal DCNN Model via Fine-Tuning and Hyperparameter Optimization

After the completion of transfer learning, one of these DCNN models that showed higher accuracy underwent further refinements through both fine-tuning and hyperparameter optimization ^59^. These refinements were made to maximize precision, to reach a prediction accuracy of as close to 100% as possible. Fine-tuning and hyperparameter optimization are iterative processes and several rounds of experimentation were required to achieve the best results for insect species identification, (cf. Figure 3). Our approach was to gradually unfreeze the last few layers of the best CNN model (that demonstrated the highest accuracy) while keeping the earlier layers frozen, Figure 6. This strategic adjustment fine-tuned the model towards learning specific features while retaining the pre-learned general representations.

**Figure 6:**
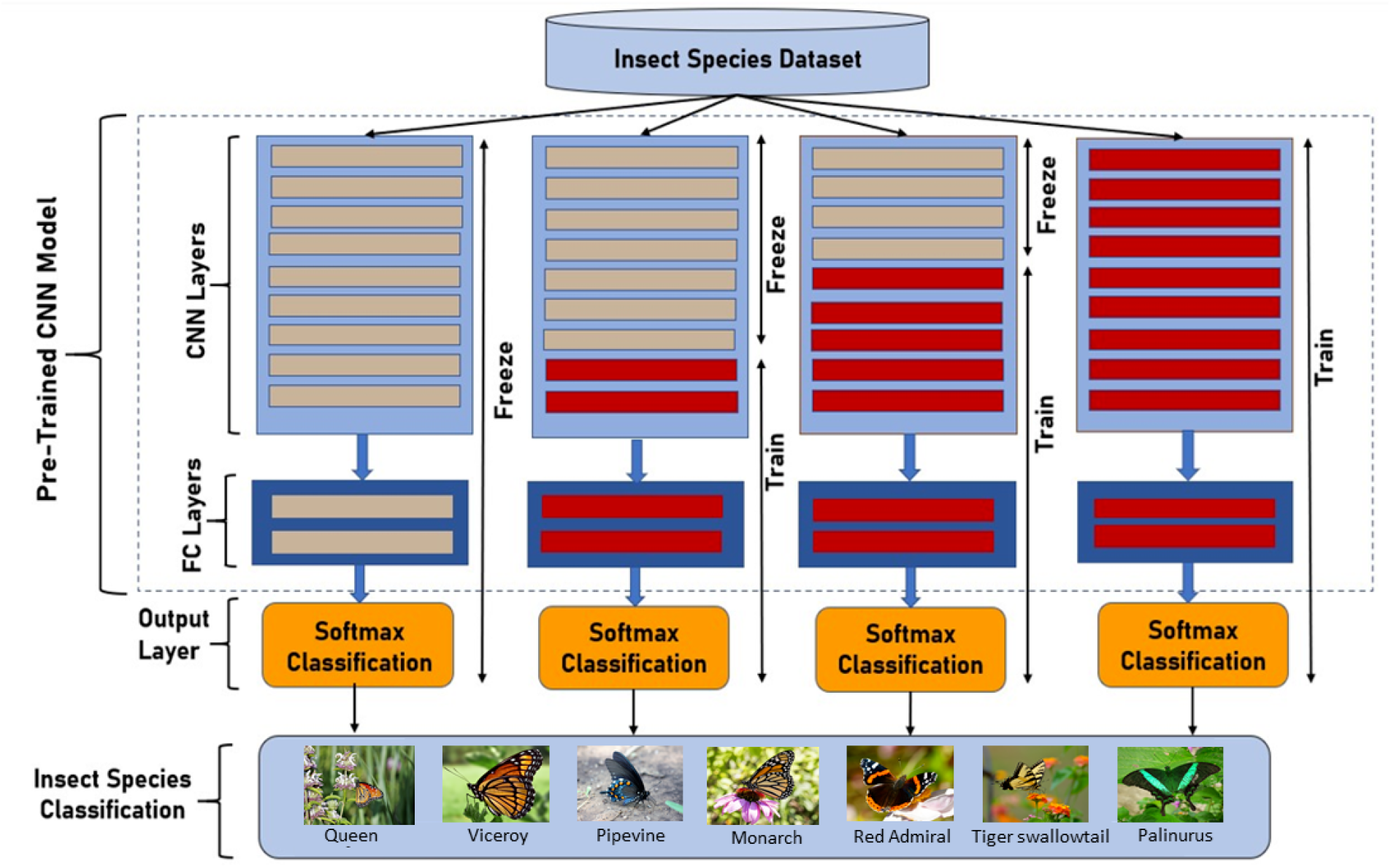
Layer Unfreezing Strategies in Fine-Tuning ResNet101: showing four strategies, progressively unfreezing the last convolutional layers. This approach helps deeper layers adapt to the new task while preserving earlier pre-trained features for effective fine-tuning.

In this work, four fine-tuning scenarios were explored to assess the impact of different configurations on model performance and adaptation to the species dataset, while also mitigating the risk of overfitting. **Overfitting** occurs when the model learns to perform well on the training set but fails to generalize to new, unseen data. To address this, we use validation accuracy and cross-entropy loss as shown in Equation 15 and Equation 3 respectively to ensure the model does not overfit. For model fitting, we used optimization algorithms to minimize loss as mentioned in the previous section. We progressively unfroze layers of the model based on performances observed in a separate validation dataset in the following sequence: the weights of the last 10 convolutional layers were first unfrozen, followed by the weights of the last 20 convolutional layers, the last 30 convolutional layers, and finally, the weights of the last 40 convolutional layers. This process involved an iterative evaluation of model performance as additional layers were unfrozen, allowing us to determine the optimal depth at which the model achieves the best performance using the insect species image dataset. During hyperparameter optimization, we systematically searched for the optimal values of hyperparameters including learning rate, batch size, dropout rate, weight decay and optimzation algorithms. These control the architecture, and training process. Hyperparameters were adjusted such that the training and validation loss would be declined when additional improvements were needed, reiterating the process to achieve these improvements. Regularly monitoring model performance on a validation set separate from the training data was conducted to ensure the accurate prediction of new, unseen data, as opposed to the mere memorization of the training set.

We used batch normalization to improve the training stability and speed of DCNN. In neural networks, the input to each layer varies significantly during training due to changes in the parameters of the preceding layers. These variations slow down training and make it more challenging to optimize the network ^23^. Batch normalization addresses the issue of input variance by normalizing the inputs to a layer within mini-batches. For each mini-batch {*x*_1_, *x*_2_, …, *x*_*m*_} of data is processed during training. Batch normalization computes the mean, *μ*_*B*_, and variance, 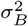, of the inputs for that mini-batch, where *m* is the batch size, as shown in Equations 10 and 11.

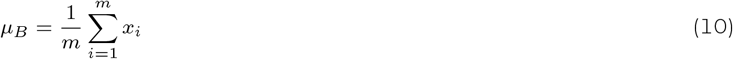

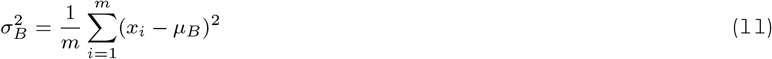

After calculating *μ*_*B*_ and 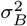, normalized activation is performed based on Equation 12, where *ϵ* is a small constant added to the variance to prevent division by zero, and 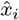 represents the normalized activation.

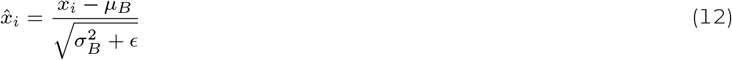

After normalization, the activation is scaled and shifted using learnable parameters *γ* and *β* as shown in Equation 13, *γ* and *β* are initialized to 1 and 0 respectively, where *y*_*i*_ represents the final normalized and transformed activation.

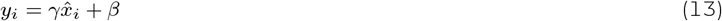

Training performance was monitored using different mini-batch sizes (8, 16, 32, and 64) with different epochs (50, 100, 150, 200). The training process minimized the loss function, *L*, by iteratively adjusting the mini-batch and epochs. Dropout is a regularization technique, that involves randomly deactivating a fraction of neurons during each forward and backward pass, thereby preventing over-reliance on specific neurons and promoting a more robust network ^25^. In this study, dropout rates ranging from 0.2 to 0.5 were systematically tested during the training process in pursuit of mitigating overfitting in the training process. Weight decay is an other regularization technique used to discourage large weights by adding a penalty term to the loss function ^23^, thereby preventing over-fitting as shown in Equation 14. Where *λ* denotes the weight decay rate, controls the strength of penalty, and *N* represents the total number of parameters in the model.

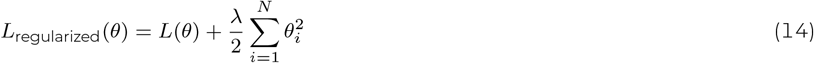

This study explored various weight decay rates ranging from 0.0001 to 0.01, to identify the optimal set of parameters. During the fine-tuning process, hyperparameters were iteratively updated to achieve optimal model performance. The objective was converge towards 100% identification accuracy, while simultaneously observing a decline in both training and validation loss. This iterative refinement of hyperparameters was important in tailoring the DL model to the specific characteristics of the insect species dataset, ensuring that it generalized effectively, prevented overfitting, promoted the convergence of training and validation accuracy, and achieved peak performance. After achieving optimal performance, the trained DL model underwent testing to evaluate its accuracy and robustness. The model was subsequently deployed as AInsectID Version 1.1 open source software, providing a user-friendly interface for insect species identification ^1^.

### Evaluation Methods

We considered accuracy the primary measure for evaluating results. The validation accuracy and loss for CNN models were assessed as shown in Equations 15 and 3, respectively. Here, *TP, TN, FP*, and *FN* indicate correctly predicted positive samples, correctly predicted negative samples, incorrectly predicted positive samples, and incorrectly predicted negative samples, respectively.

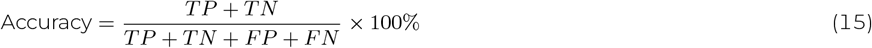

Gradient-weighted class activation mapping (Grad-CAM) identifies the specific regions of the insect mimics that contribute most significantly to the model predictions. It calculates the weights *α*^*k*^ of the feature map using global average pooling as shown in Equation 17, where, *z* is the total number of spatial locations in the feature map. This weight represents the importance of each feature map channel for predicting class *k*. The Grad-CAM heatmap *L*^*k*^ is generated by taking a weighted sum of the feature maps as shown in Equation 16. The ReLU function ensures that only positive influences contribute to the final heatmap, highlighting the areas most relevant to the prediction of class *k*.

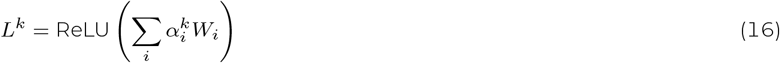

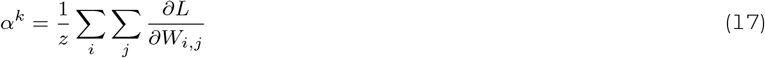

### Software Setup

MATLAB (2023a) software was used to train our CNN model. The flexibility and user-friendly nature of MATLAB facilitated seamless experimentation, enabling us to focus on model architecture and hyperparameter tuning. To harness the computational efficiency required for training large-scale CNN, we leveraged a high-performance computing environment equipped with a state-of-the-art NVIDIA GPU. The AInsectID Version 1.1 GUI software ^1^ was developed using the MATLAB App Designer Toolbox.

## RESULTS AND DISCUSSION

### Feature Representation Dynamics during Transfer Learning

Figure 7 presents the least significant and most significant feature maps across different layers of the various deep learning models, VGG16, GoogLeNet, MobileNetV2, InceptionNetV2, ResNet50, and ResNet101, that have been used for insect identification. In each model, the least significant feature maps correspond to regions where the network assigns minimal importance, often reflecting non-discriminative areas such as background or parts of the insect that are less relevant for classification. These maps are characterized by low activation values and weak gradient responses during backpropagation, indicating their limited contribution to the prediction. Conversely, the most significant feature maps highlight the areas that the networks focus on to make accurate predictions. These maps show strong activations and higher gradient responses, signifying their critical role in the classification process.

**Figure 7:**
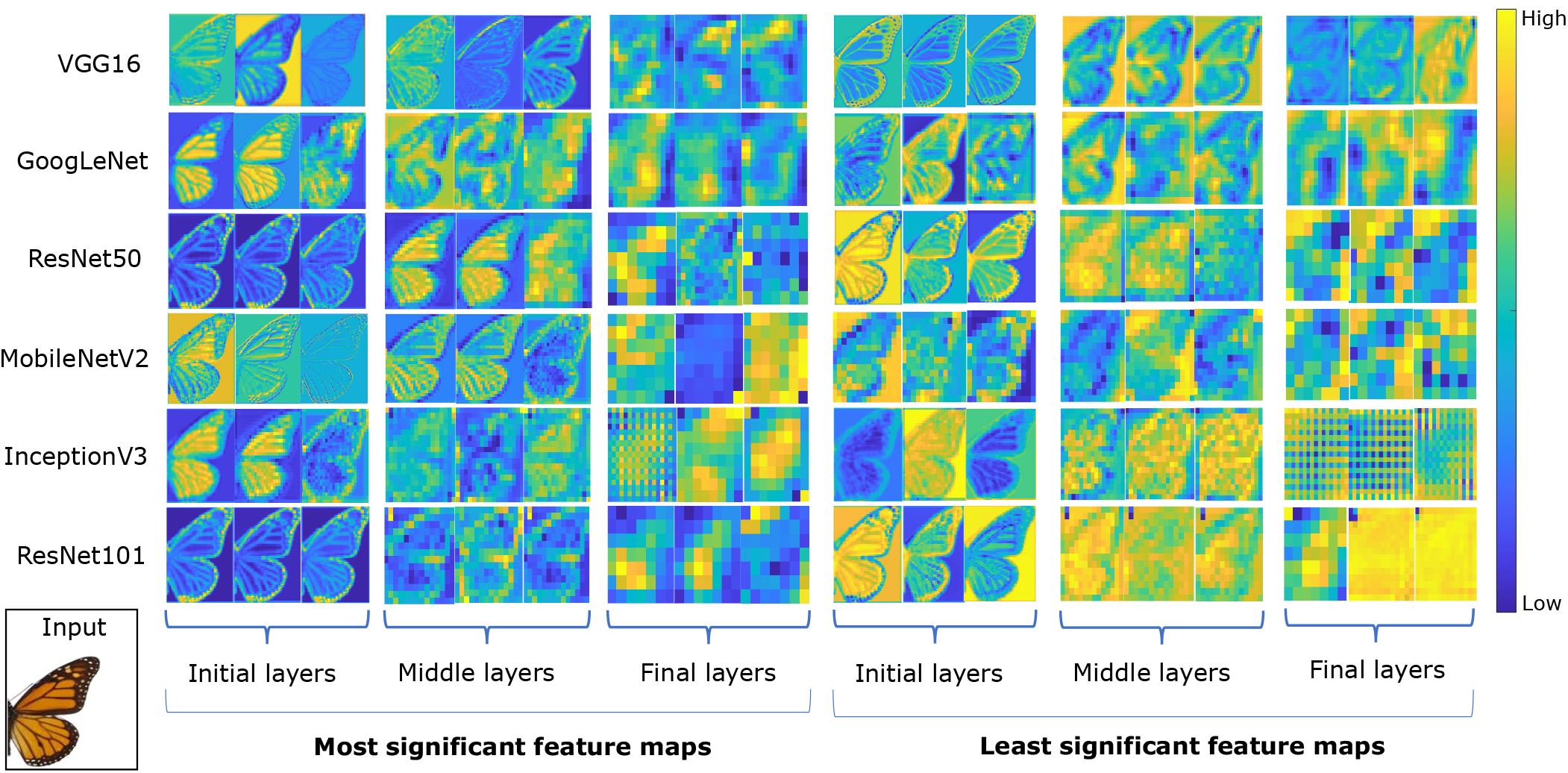
Comparison of least significant and most significant feature maps across different convolutional layers (low, middle, and high) for multiple CNN networks. The images demonstrate how feature map activations evolve, highlighting differences in the focus of the networks at different depths

The most significant feature maps in deeper architectures like ResNet101, observably capture more abstract and detailed features, while VGG16 architectures focus on simpler patterns within earlier layers. GoogLeNet, MobileNetV2, and inceptionV3 strike a balance between efficiency and depth, showing clear differences in how they handle low- and high-level features. ResNet50 and ResNet101, with their deeper residual connections, exhibit more refined feature maps in higher layers, capturing the complex textures and subtle distinctions of insect morphology

### Transfer Learning Results with and without Data Augmentation

As shown in Table 1, The use of data augmentation leads to significant improvements in the validation accuracy of all CNN models tested. For example, the accuracy of VGG16 increased from 30% to 45%, while deeper models like ResNet50 and ResNet101 saw substantial gains, improving from 78% to 88% and 85% to 92%, respectively. These results demonstrate the effectiveness of data augmentation in enhancing model performance, particularly in more complex architectures. Overall, data augmentation improves validation accuracy, helping the models generalize more effectively whilst mitigating overfitting. The dataset without data augmentation consists of 40,000 samples, while with data augmentation, the sample size increases to 61,000. The extensive depth of ResNet101, featuring 101 layers with a total 347 layers, contributes to its capacity for the learning of intricate hierarchical representations. However, this depth also requires a more extensive parameter space and increased computational demands during the training process. As a result, the training time for ResNet101 was higher that the other CNN architectures.

**Table 1:**
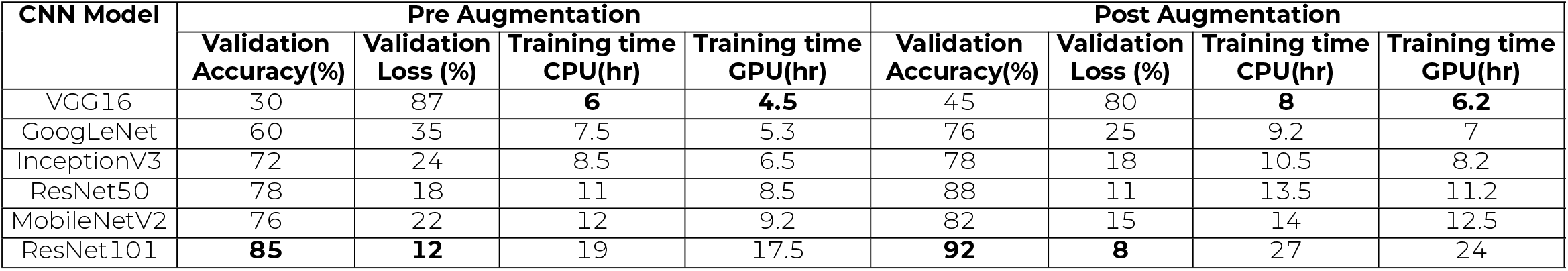
Effectiveness of data augmentation in enhancing model performance across different CNN architectures by comparing the validation accuracy and validation loss with and without data augmentation. The pre-augmentation phase consists of 410,000 images, while the post-augmentation phase contains 61,000 samples. We also present a training time comparison between CPU and GPU implementations for the same CNN models, highlighting differences in computational efficiency. The optimal results in each column are emboldened.

Figure 8 illustrates the validation accuracies and corresponding losses achieved using different learning rates, while employing various optimizers SGD, SGDm, Adam, and RMSprop. VGG16 recorded the lowest validation accuracy at 45%, likely due to its simpler architecture struggling with the complex visual patterns of the dataset. GoogleNet showed improvement with a validation accuracy of 76%, attributed to its inception modules capturing information at various scales. InceptionV3, an evolution of GoogleNet, further increased the accuracy to 78% through more efficient convolution factorizations. MobileNetV2 achieved a validation accuracy of 82%. Notably, ResNet50 achieved the validation accuracy at 88%, benefitting from its deep residual learning framework that effectively mitigates the vanishing gradient problem, enabling better performance in detailed and variable visual data recognition. However ResNet101 emerged as the most effective model, outperforming other well-known architectures. The depth of ResNet101, which includes 101 layers, proved crucial in capturing the detailed and subtle features needed for accurately classifying diverse insect species with 92% accuracy. The achieved 92 % validation accuracy not only signifies the model’s robustness but also positions ResNet101 as a potent candidate for fine-tuning, where precision and accuracy are paramount.

**Figure 8:**
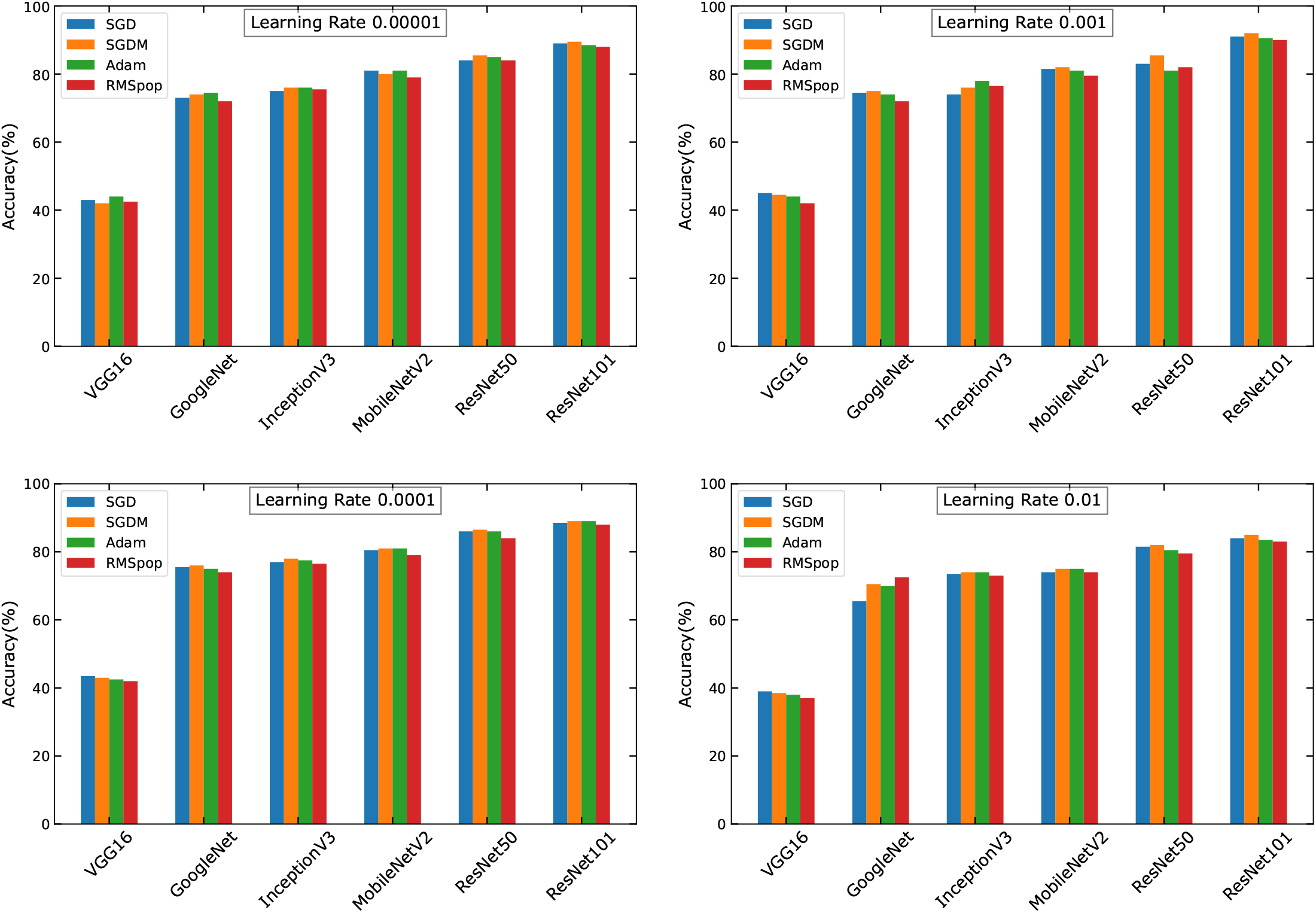
Comparison of optimization algorithms’ accuracy illustrates the performance comparison of SGD, SGDM, Adam, and RMSProp optimization algorithms in terms of accuracy across CNN models to evaluate how each optimizer affects the overall accuracy of the models.

As shown in Figure 8, minor differences were observed among the accuracies of SGD, SGDM, Adam, and RMSprop optimizers with different learning rates. However SGDM demonstrated slightly higher accuracy of 92% with a learning rate of 0.001 as compare to others. This suggests that the incorporation of momentum in the optimization process contributed to improved performance, albeit marginally, compared to other commonly used optimizers. In Figure 9, we investigate the impact of varying batch sizes across different learning rates on the performance of CNN models. A batch size of 32 emerges as the optimal choice across all CNN models, consistently demonstrating the highest accuracy across different learning rates in our study. For ResNet101, the combination of a batch size of 32 with a learning rate of 0.001 resulted in the highest accuracy compared to other configurations. In this study, each architecture was trained for a maximum of 150 epochs, we observed that the training process encountered local minima, where the loss does not improve significantly, resulting in a steady accuracy level beyond 100 iterations as shown in Figure 10. Despite continuing the training process, further improvements in accuracy were not achieved, suggesting that the models had reached a plateau in learning capability within 100 iterations.

**Figure 9:**
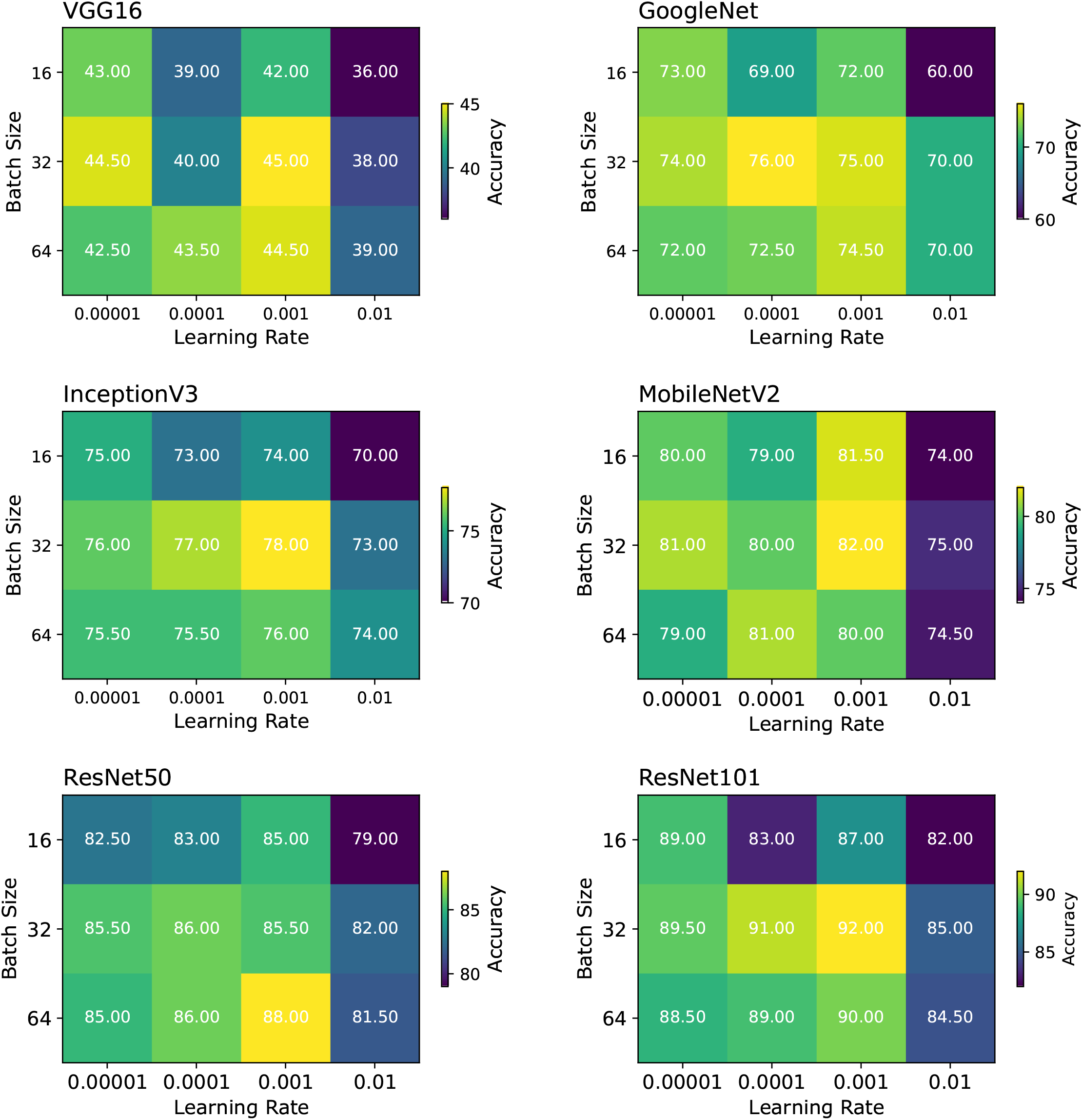
The choropleth maps illustrate the impact of batch size and learning rate on the performance of various CNN architectures, visualizing how changes in these hyperparameters influence model accuracy and efficiency

**Figure 10:**
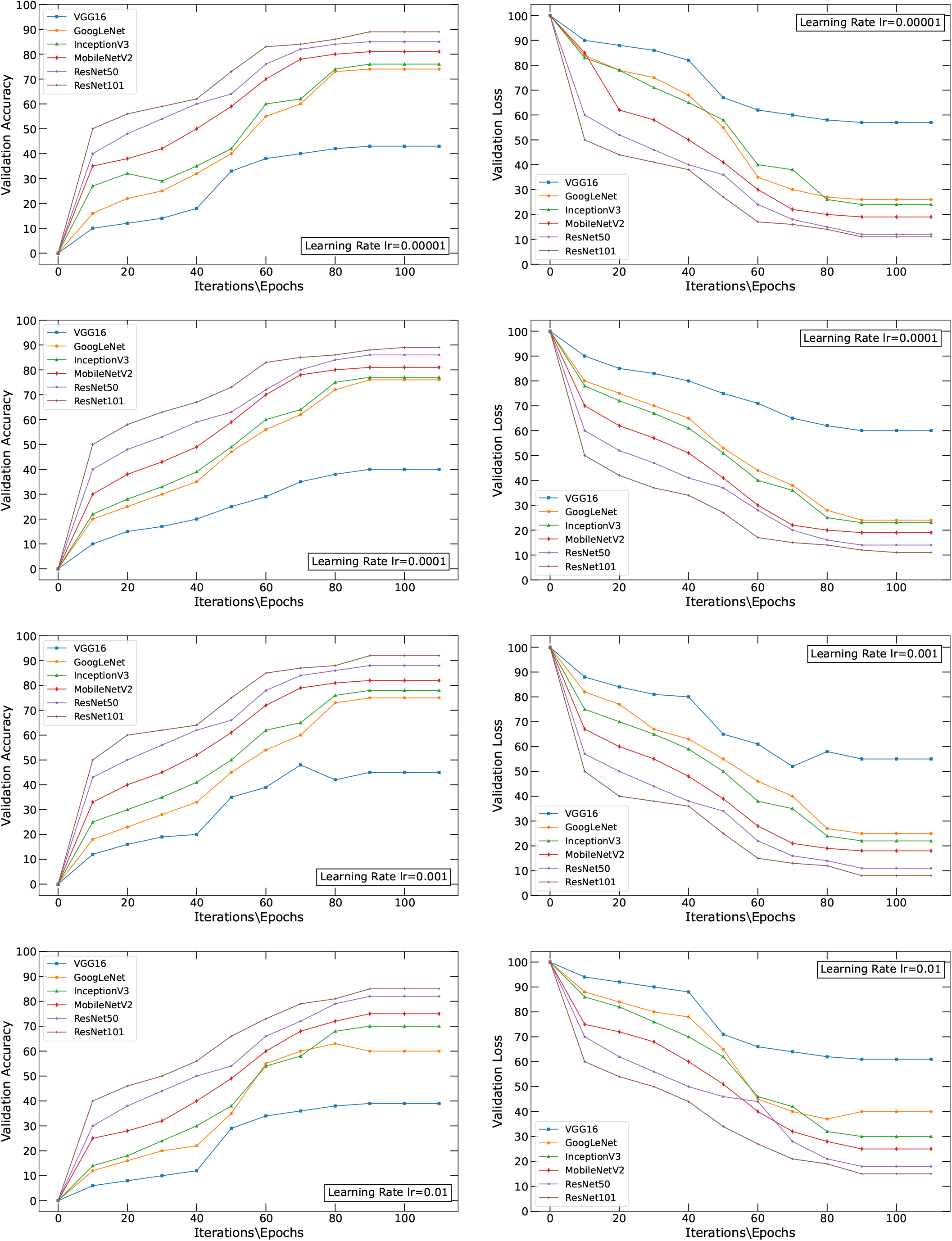
Comparison of Highest Accuracy and Loss Across CNN architectures with different Learning Rates showcasing the highest achieved accuracy and corresponding loss metrics across the CNN architectures. Each architecture was trained for a maximum of 150 epochs, with results indicating the epoch at which local minima were reached.

### Hyperparameter Optimization and Fine Tuning of ResNet101

Based on the transfer learning results, the decision to focus on ResNet101 for fine-tuning was made due to its exceptional performance compared to the other models considered such as VGG16, GoogLeNet, ResNet50, MobileNetV2, and InceptionV3. In our experimentation with fine-tuning a ResNet-101 model by systematically varying the number of unfrozen layers to investigate its impact on model performance as shown in Table 2. The results demonstrate the impact of gradually unfreezing layers during the fine-tuning process. Starting with a conservative approach of unfreezing only 10 layers, the model achieves moderate improvements in both training and validation accuracy compared to the initial frozen model. This gradual unfreezing strategy allows the model to adapt to the specific features of the dataset while minimizing the risk of overfitting. Continuing to unfreeze additional layers in increments of 10 (Unfreeze 20 Layers, Unfreeze 30 Layers) results in further enhancements in accuracy, culminating in a remarkable training accuracy of 100% and a validation accuracy of 99.65% after unfreezing 30 layers. This progressive unfreezing approach enables the model to leverage deeper layers to learn more intricate features in the data, leading to improved performance. Nevertheless, after unfreezing more than 30 layers (to a maximum of 40 Layers), the accuracy begins to decline. This decline may be attributable to the increasing complexity of the model and the risk of overfitting. Moreover the results reveal a notable relationship between the number of trainable parameters and the processing time required for training the model. An increase in the number of trainable parameters augments model complexity, the cost for which is computational time.

**Table 2:** The effect of gradually unfreezing layers on model performance, showing improvements in validation accuracy and reductions in validation loss as more layers are fine-tuned, while also highlighting reductions in training time when utilizing GPU acceleration compared to CPU. Superior values in training accuracy, validation accuracy and validation loss are emboldened.

| Numbers of Fine Tune Layers | Training Accuracy | Validation Accuracy | Validation Loss | Trainable Parameters | CPU Time (Hours) | GPU Time (Hour) |
| --- | --- | --- | --- | --- | --- | --- |
| Unfreeze 10 Layers | 96% | 94.60% | 0.28% | 4465664 | 38.50 | 26.20 |
| Unfreeze 20 Layers | 98% | 97.20% | 0.16% | 8931328 | 40.25 | 27.50 |
| Unfreeze 30 Layers | **100%** | **99.65%** | **0.03%** | 14450176 | 44.20 | 29.50 |
| Unfreeze 40 Layers | 99% | 98% | 0.09% | 15831308 | 46.30 | 31.40 |

The choropleth maps in Figure 11 reveal how accuracy changes as different numbers of layers are unfrozen during training. Each choropleth map shows a specific set of unfrozen layers, with the range being between 10 and 40, along with different batch sizes and learning rates. Unfreezing 10 layers shows moderate accuracy improvements, with optimal performance observed at a learning rate of 0.0001 and a batch size of 32. When 20 layers are unfrozen, further enhancements can be observed, consistently nevertheless, favoring a learning rate of 0.0001. When 30 layers are unfrozen, significant improvements in accuracy are observed, particularly with 32 batch sizes and at a learning rate of 0.0001. Conversely, the validation loss demonstrates a consistent decrease as the number of fine-tuned layers is raised. The lowest validation loss of 0.03% is attained when 30 layers are unfrozen. However, with 40 unfrozen layers, accuracy improvements were notably marginal, indicating diminishing returns as the layer was count increased.

**Figure 11:**
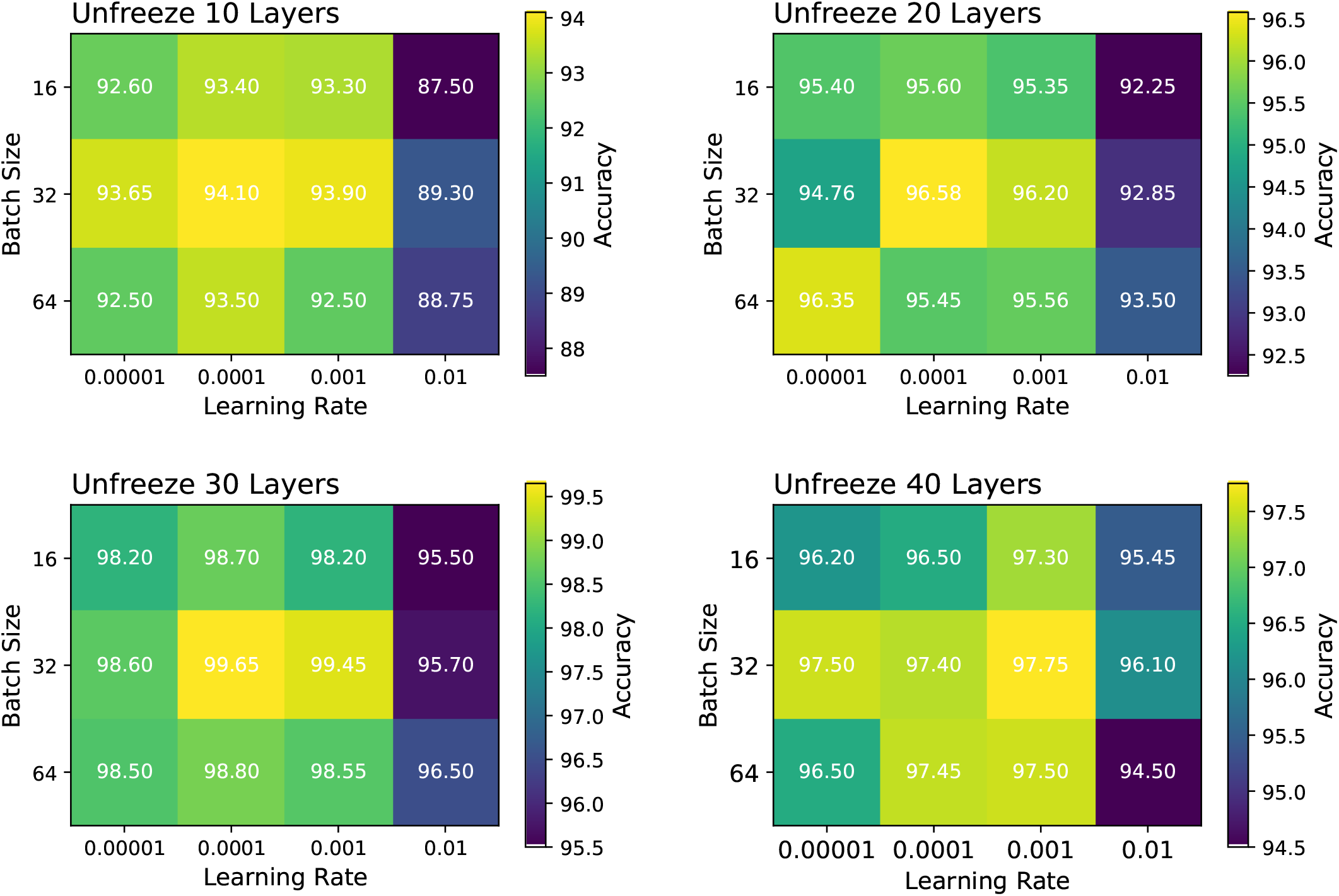
The choropleth maps illustrate the impact of batch size and learning rate on the fine-tuning results after progressively unfreezing layers of ResNet101

Based on these results the final fine-tuning of ResNet101 involved SGDM optimization with a learning rate of 0.0001, a minibatch size of 32, and regularization through dropout (0.3) and weight decay (0.001), to prevent overfitting. By unfreezing the last 30 layers, the model was able to adapt to the specific dataset while retaining the general features learned during pre-training. GPU acceleration was utilized to expedite the training process, ensuring efficient convergence. This combination of hyperparameters, as shown in Table 3 resulted in optimal performance, with the model achieving 100% training accuracy and 99.65% validation accuracy with robust generalizations for insect species dataset including mimics.

**Table 3:** Optimal hyperparameter settings identified for fine-tuning ResNet101, which resulted in the highest validation accuracy.

| Hyperparameters Options | Optimal Values |
| --- | --- |
| Optimization Algorithms | SGDM |
| Learning Rate | 0.0001 |
| Minibatch Size | 32 |
| Momentum | 0.9 |
| Maximum Epochs | 100 |
| Dropout Rate | 0.3 |
| Dropout period | 5 |
| Weight Decay | 0.001 |
| Execution Environment | GPU |

As shown in Figure 12, the gradient-weighted class activation map (Grad-CAM) results for the fine-tuned ResNet101 model provide valuable insights into its interpretation of images of the three mimic models including *Danaus plexippus, Delias Belisama*, and *Battus philenor*. Each choropleth map highlights the regions of the images that the model deemed most significant when classifying them into their respective categories. Areas with higher activation are indicated in red, illustrating the model’s focus on specific features pertinent to its classification task. This visualization underscores the ability of the model to leverage nuanced details in the images, revealing the underlying features that influence its predictions. By analyzing these maps, the fine-tuned ResNet101 model prioritization of certain regions can be discerned, enhancing its decision-making process in a complex classification environment.

**Figure 12:**
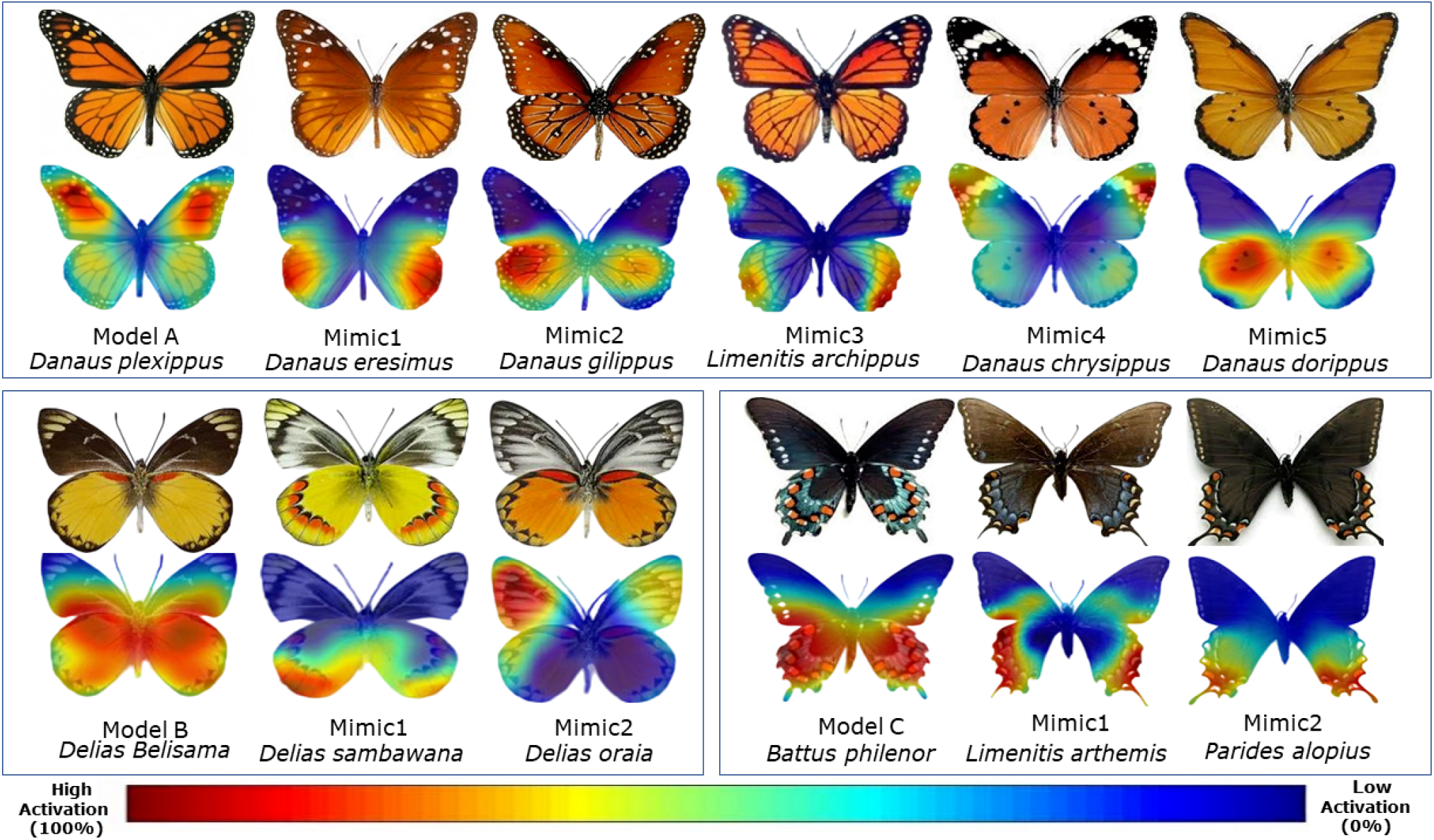
Results of the fine-tuned ResNet101 gradient-weighted class activation map (Grad-CAM). These provide insights into how the finetuned ResNet101 model interprets images from each of the three mimic models. These choropleth maps show the regions of the images that were most important for model classification of images into respective categories.

In Figure 13, box-and-whisker plots are used to show the accuracies of the fine-tuned ResNet101 model across the three different mimic datasets, with 25 test images used for each species. A one-way ANOVA test was used to evaluate the levels of statistical difference in terms of predictive performance across the three mimic models. Using an *α* value of 0.05, we find that all the p-values from the one-way ANOVA test are above 0.05. This indicates that there are no statistical differences in predictive accuracy of the ResNet101 model across the three mimic datasets. This means that the fine-tuned ResNet101 model performs similarly across all three mimic species (*Danaus plexippus, Delias belisama*, and *Battus philenor*). The higher p-values suggest that any observed differences in accuracy between the datasets are likely due to random variation, rather than any meaningful difference in model performance. This further implies that the model is consistently able to classify images across the different mimic models.

**Figure 13:**
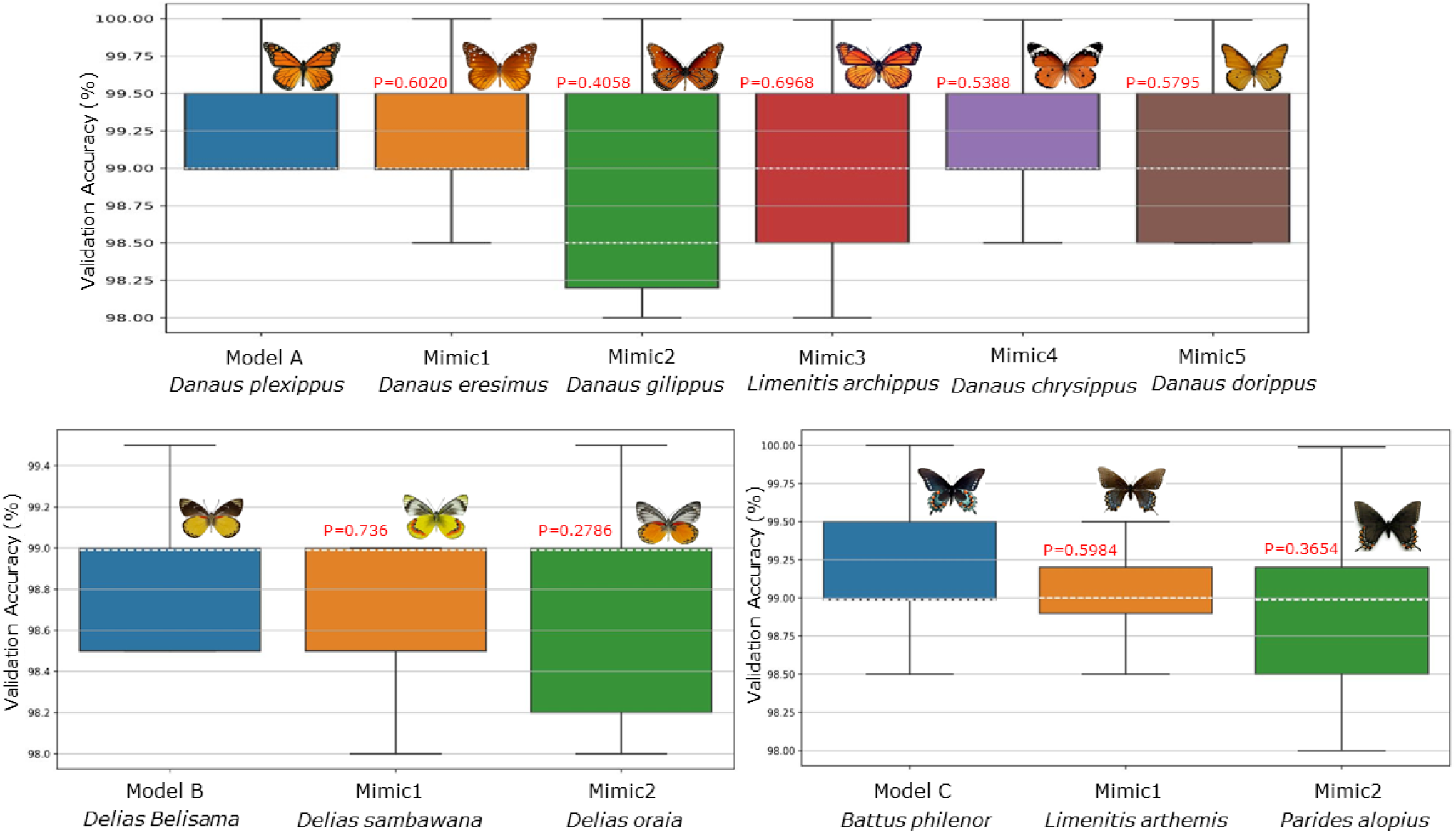
Box-and-whisker plots showing the accuracies of the ResNet101 model across three different mimics models, with 25 test images used for each species in the representation (n = 25). The whiskers extend to the minimum and maximum accuracy values, excluding outliers. A one-way ANOVA test using *α* = 0.05 was conducted to evaluate the statistical significance of the differences in fine-tuned ResNet101 model performance across the datasets. A resulting p-value *<* 0.005 indicates a statistically significant difference among accuracies, and *p* ≥ 0.05 indicates there is no statistically significant difference. In every case, we find *p* ≥ 0.05.

## Disscusion

The research presented here, is concerned with the identification of insect mimics utilizing a suite of advanced deep learning architectures including VGG16, GoogLeNet, InceptionV3, MobileNetV2, ResNet50, and ResNet101. The integration of dataset augmentation techniques further enhances model generalization across various types of insect mimicry. Among these models, ResNet101 has emerged as the most effective, demonstrating the highest accuracy in identifying insect mimics after the application of transfer learning techniques. Our analyses present the nuanced impacts of different optimization algorithms and hyperparameters on model performance. Subtle variations in accuracy among optimizers like SGD, SGDM, Adam, and RMSprop, particularly at different learning rates, are notable. SGDM stands out marginally, showcasing the influence of momentum incorporation. The momentum of the SGDM component aided in smoother convergence through the accumulation of velocity across iterations (cf. Figure 8), which were important for navigating complex optimization landscapes. Finding the appropriate learning rate was crucial. If the learning rate was too high (0.01), the optimization process diverged, leading to unstable training or overshooting the optimal solution (cf. Figure 9).

When fine tuning of Resnet101, the training and validation accuracies exhibited a discernible pattern (cf. Table 2), each generally improving, and reaching a peak of 100% training accuracy and 99.65% validation accuracy on unfreezing 30 layers. Table 4 compares this study against other recent works on insect mimic identification. Specifically, the table compares the accuracy, learning rate, otpimizer used, batch size and CNN model in each case. With an accuracy of 99.65%, our results outperform the limited number of published studies on insect mimic identification, highlighting the effectiveness of the proposed approach in a relatively underresearched area.

**Table 4:** Comparison of accuracy results from recent published works for insect mimic identification. The table lists the accuracy, learning rate, optimizer, batch size, CNN models, and fine tuning approaches for each referenced publication.

| Publication | Year | Accuracy | Learning rate | Optimizer | Batch size | CNN model | Fine tuning method |
| --- | --- | --- | --- | --- | --- | --- | --- |
| Wang et al.<sup>60</sup> | 2022 | 96% | 0.0001 | Adam | 64 | S-ResNet | Residual structure |
| Kalaiarasi et al.<sup>61</sup> | 2023 | 88.56% | 0.0001 | Nadam | 32 | InceptionV3 | Softmax layer |
| Doan et al.<sup>62</sup> | 2023 | 72.31% | 0.001 | Adam | 32 | EfficientNet | Power Mean SVM |
| Hassan et al.<sup>63</sup> | 2024 | 88.48% | 0.001 | Adam | 32 | ResNet101-SA | Self attention layers |
| Ong et al.<sup>64</sup> | 2024 | 89% | 0.00001 | Adam | 79.5 | Xception | Convolutional blocks |
| Mattins et al.<sup>65</sup> | 2024 | 98.20% | 0.001 | SGD | 64 | Effi-CNN-3 | Convolutional blocks |
| Hasan et al.<sup>66</sup> | 2024 | 97% | 0.0001 | SGD | 32 | ResNet152V2 | Unfreeze all layers |
| Spiesman et al.<sup>67</sup> | 2024 | 94.9% | 0.001 | Adam | 32 | EfficientNetV2 L | Freeze all layers |
| Manchanda et al.<sup>68</sup> | 2024 | 83.99% | 0.001 | Adam | 32 | Xception | Unfreeze all layers |
| Sauer et al.<sup>69</sup> | 2024 | 91% | 0.0001 | Adam | 32 | ResNet101 | Unfreeze all layers |
| **This study** | 2024 | **99.65%** | 0.0001 | SGDM | 32 | ResNet101 | Unfreeze last 30 layers |

Conversely, the validation loss demonstrates a consistent decreasing trend as the number of fine-tuned layers increases. The lowest validation loss of 0.03% is attained when 30 layers are unfrozen. This indicates that deeper fine-tuning allows the model to capture finer details and nuances in insect images, leading to improved generalization performance and more accurate species identification. Notably, our experiments revealed that while varying batch sizes influenced model performance across the insect dataset, the most optimal results were consistently achieved with a batch size of 32 with learning rate 0.0001.

Our comprehensive tuning strategy proved the best fit to strike a delicate balance between model complexity and robustness, ultimately seeking improved accuracy and generalization, leveraging the parallel processing capabilities of a GPU environment. Gradually unfreezing layers effectively improved the performance of the model, indicating a successful fine-tuning strategy, such that the model learned intricate features within the insect dataset without sacrificing its ability to generalize to unseen examples.

Table 4 provides a comparative overview of CNN-based models for insect species identification, revealing a range of accuracies reported in recently published articles. Models like ResNet variants, EfficientNet, InceptionV3, and custom architectures (e.g., Effi-CNN-3) exhibit varying levels of success, with ResNet-based models proving to be consistently high in performance, this being due to features like self-attention and residual structures. Fine-tuning strategies differ widely. Some studies, such as Wang et al. ^60^, use residual layer modifications, while others (e.g., Manchanda et al. ^68^) unfreeze all layers, impacting the prediction accuracy. Our study selectively unfreezes the last 30 layers of ResNet101, and combined with SGDM optimization at a learning rate of 0.0001, achieves the highest prediction accuracy compared to any other recent study at 99.65%. This suggests that targeted fine-tuning to preserve feature quality, while concurrently enhancing specificity, is an effective approach. Adam has been a preferred optimizer in the majority of recent studies, though SGDM as used in our study, has proved beneficial for achieving precise training outcomes. Variations in batch size, optimizer, and learning rate are noteed to affect model performance, which depend to a great extent on finely tuned hyperparameters and careful architectural adjustments, highlighting our success in selective tuning and optimization to enhance classification accuracy.

## CONCLUSIONS

AInsectID v1.1 open source software represents an advancement in the field of insect identification, achieving an accuracy rate of 99.65%, and outperforming the accuracy rates from previous studies. This study addresses challenges related to the identification of insect species using Deep Convolutional Neural Networks (DCNNs), with a focus on leveraging advanced transfer learning and fine-tuning strategies to enhance identification performance. Our research problem was primarily concerned with achieving high accuracy in distinguishing between insect species, and testing the prediction accuracy against insect mimics. By evaluating several CNN models and comparing their accuracies, fine-tuning approaches, and performances across key parameters, we provide a comprehensive exploration highlighting the potential of CNN in insect species classification. The main contributions of our work are in the identification of optimal model structures and hyperparameters for insects mimics. After rigorous testing, we found that ResNet101, together with the selective unfreezing of the last 30 layers, offers a high level of balance between generalization and specificity, achieving a peak accuracy of 99.65%. Using a selective fine-tuning approach, coupled with SGDM optimization and an appropriate learning rate, was found to significantly enhance model performance, without the computational burden of training from scratch. When comparing against other published studies using different CNN models and fine-tuning strategies, our approach demonstrates that by preserving essential feature extraction layers while adjusting recent layers, we are able to optimally leverage transfer learning through challenging classification tasks. Beyond model selection and hyperparameter tuning, our research elucidates the broader applicability of fine-tuned CNNs in detailed species identification tasks. By achieving nearly perfect classification accuracy, AInsectID v1.1 offers promising implications for expanding DCNN use in biodiversity and conservation fields. Additionally, our work shows that CNNs can handle complex visual tasks, such as the differentiation of mimic species from their mimicked counterparts, an accomplishment that could extend to identifying subtle phenotypic differences in other taxa as well. There remain nevertheless several avenues for future directions for this work. Expanding the dataset to include more mimic species could be used to test model robustness, as well as its ability to generalize to unfamiliar species. Research into automated data augmentation, using generative models, might also yield richer training data, further boosting classification accuracy.

## DATA AVAILABILITY

AInsectID v1.1 is available as an open source package from https://doi.org/10.7488/ds/7801. The package includes all Matlab codes, as well as the Graphical User Interfaces for the software. The training datasets used in this work are owned and stored by the National Museum of Scotland, Edinburgh, and the Natural History Museum, London. The Zenodo (CERN Data Center) open source dataset is available at: https://doi.org/10.5281/zenodo.3549369

## ACKNOWLEDGMENTS

The authors wish to thank Dr. Marcelo Dias from The University of Edinburgh for his helpful advice and productive comments. H.S. wishes to express appreciation to the Higher Education Commission (HEC) of Pakistan for providing a fully funded Ph.D. scholarship. All authors are grateful to the National Museum of Scotland and the Natural History Museum, London, for generously providing the insect species datasets used in this study.

## AUTHOR CONTRIBUTIONS

Conceptualization: P.A. Methodology: H.S. and P.A. Investigation: H.S. and P.A. Visualization: H.S. Writing: H.S. Editing: P.A. Funding Acquisition: H.S. Supervision: P.A.

